# The *dgc2* gene encoding di-guanylate cyclase suppresses both motility and biofilm formation in the filamentous cyanobacterium *Leptolyngbya boryana*

**DOI:** 10.1101/2022.01.10.475764

**Authors:** Kazuma Toida, Wakana Kushida, Hiroki Yamamoto, Kyoka Yamamoto, Kazuma Uesaka, Kunio Ihara, Yuichi Fujita, Hideo Iwasaki

## Abstract

Colony pattern formations of bacteria with motility manifest complicated morphological self-organization phenomena. *Leptolyngbya boryana* is the filamentous cyanobacterial species, which has been used as a genetic model organism for studying metabolism including photosynthesis and nitrogen-fixation. Although a widely used type strain (wild type) of this species has not been reported to show any motile activity, we isolated a spontaneous mutant strain which shows active motility (gliding activity) to give rise to complicated colony patters, including comet-like wandering clusters and disk-like rotating vortices on solid media. Whole-genome resequencing identified multiple mutations on the genome in the mutant strain. We confirmed that inactivation of a candidate gene, *dgc2* (*LBDG_02920*), in the wild type background was sufficient to give rise to motility and the morphological colony patterns. This gene encodes a protein, containing the GGDEF motif, which is conserved at the catalytic domain of diguanylate cyclase (DGC). Although DGC has been reported to be involved in biofilm formation, the mutant strain lacking *dgc2* significantly facilitated biofilm formation, suggesting a role of DGC for suppressing both gliding motility and biofilm formation. Thus, *L. boryana* provides an excellent genetic model to study dynamic colony pattern formation, and novel insight on a role of c-di-GMP for biofilm formation.

**Importance:** Self-propelled bacteria often show complicated collective behaviors, such as the formation of dense moving clusters, which is exemplified by wandering comet-like and rotating disk-like colonies, while molecular details of forming such structures remain limited. We found that a strain deficient in the diguanylate cyclase gene *dgc2* in the filamentous cyanobacterium *L. boryana* induces motility, complex and dynamic colony pattern formation including the comet-like and disk-like clusters. While c-di-GMP has been reported to activate biofilm formation in some bacterial species, disruption of the *dgc2* gene unexpectedly enhanced it, providing a novel role for c-di-GMP regulatory system in both colony pattern formation and biofilm formation.

## Introduction

Many bacterial species exhibit morphological colony pattern formations with motility. For example, *Paenibacillus vortex* and *Pseudomonas aeruginosa* shows complex colony formation with swarming motility (1, 2 2). Mechanisms underlying motility and morphological formation, and environmental impacts on them have been studied, and theoretical analysis has also provided some insights on the self-organization processes (3-6;). In cyanobacteria, which lack flagella, twitching and gliding motility have been studied with some species, including *Oscillatoria* (7, 8), *Phormidium* (4), hormogonia of *Nostoc punctiforme* (9, 10) and phototactic movement of the unicellular *Synechocystis* sp. PCC 6803 (hereafter, *Synechocystis*, 11, 12). Twitching motility indicates crawling on the surface using the extension and contraction of the Type IV-pilus (13). Gliding of filamentous cyanobacteria is suggested to be driven by a polysaccharide secretion system, the junctional pore complex (JPC). In hormogonia of *N. punctiforme*, JPC is formed with arrayed ring structure of Type IV-pilus-like systems, which are in part consist of Pil and Hps proteins (9, 14-16). The arrayed ring-accompanying JPC has also been observed in *Phormidium* species which forms spiral clusters (4; 17). We recently reported that the filamentous cyanobacterium, *Pseudanabaena* sp. NIES-4403 (hereafter, *Pseudanabaena*) generates high-density migrating clusters, which were categorized into comet-like wandering and disk-like rotating clusters (18). The combination of these wandering and rotating clusters has been reported in *Bacillus* (19, 20;), *Paenibacillus vortex* in the presence of mitomycin C (21), and in *Paenibacillus* sp. NAIST15-1 (22). Our previous observations suggested in *Pseudanabaena* that the following processes are key to generate the wandering and rotating clusters: (1) follow-up motion of bacterial filaments via polysaccharide secretion or groove formation on the solid-phase surface, (2) formation of bundles with nematic alignment, (3) formation of comet-like clusters accompanied by the creation of a cover of filaments at the tip in the direction of travelling, (4) formation of rotating disks by spontaneous self-tracking of comet-like clusters, and (5) transition based on collisions between different types of clusters (18). Nevertheless, molecular details of generating these colony patterns remain largely unknown: genetic study has been limited to *Paenibacillus* sp. NAIST15-1 (22), where a large extracellular protein, CmoA was identified to facilitate motility and migrating cluster formation.

*Leptolyngbya boryana* (*L. boryana*, hereafter), previously known as *Plectonema boryanum*, is a non-heterocystous, filamentous cyanobacterium belonging to Section III (23). This species is able to fix nitrogen under microoxic conditions (24) and to heterotrophically grow in the presence of glucose in the dark (25). Since it is genetically tractable, *L. boryana* provides an excellent model to study photosynthesis and nitrogen fixation (26-28). The *dg5* strain, referred to as a wild type strain in the present study, is a variant which is able to heterotrophically grow under complete darkness in the presence of glucose (27). Although this strain has not been reported to show active motility, we isolated a spontaneous mutant strain, named E22m1’, which showed active gliding motility and colony morphology with a mixture of comet-like wandering clusters and rotating clusters on the surface of solid media. Comparison of genomic DNA sequences between the wild type and E22m1’ strains led to identify multiple mutations. We found disruption of a gene named *dgc2* encoding cyclic di-GMP (c-di-GMP) synthetase (diguanylate cyclase, or DGC) activates the gliding motility. DGC catalyzes synthesis of c-di-GMP, an intracellular signaling molecule, from two molecules of GTP. The catalytic site of DGC consists of characteristic amino acid sequence motif, GGDEF or GGEEF (29), referred to as the GGDEF motif. c-di-GMP-dependent inhibition of cell motility has been reported in many bacterial species, such as *Escherichia coli, Pseudomonas aeruginosa* and *Acinetobacter baumannii* (30-32). In cyanobacteria, Cph2 is known as a DGC in *Synechococcus* for c-di-GMP production, which inhibits pili-based motility (33, 34). Cph2 harbors a photoreceptive GAF domain which controls phototaxis in *Synechocystis* (34). Moreover, c-di-GMP has been reported to facilitate biofilm formation in many bacterial species (35, 36); in the thermophilic cyanobacterium, *Thermosynechococcus vulcanus*, c-di-GMP directly binds to a cellulose synthase to facilitate cellulose synthesis and cell aggregation (37, 38).

## Results and Discussion

### Nullification of a diguanylate cyclase gave rise to motility in *L. boryana*

The wild type strain of *L. boryana* does not show gliding motility (**Movie S1**, upper left panel). When cell suspension was inoculated at the center of solid media on a 90-mm dish, most of grown cells remaining at the inoculation spot. Within the colonies, the wild type filaments intertwined with each other, forming dense knot-like clumps locally. At the edges some elongated filaments protruded, giving an intertwined ivy-like morphological pattern (**Fig. 1A**, left panel). By contrast, we have isolated a spontaneous mutant, tentatively named E22m1’, which turned to move on agar plate (**Fig.1A**, middle left; **Movie S1**, upper right panel). More precisely, E22m1’ was isolated as a spontaneous mutant from a non-motile, *dg5*-derived strain, E22m1-*dg5*. The E22m1-*dg5* is a transformant in which a kanamycin-resistance gene had been inserted into the region between ORFs *LBDG_21990* and *LBDG_22000*, which did not show any phenotypic changes. After long-term maintenance of E22m1-*dg5* on BG-11 agar plates for about 1.5 years, we found a subpopulation of cells showed motility on an agar plate. After picking up the motile colonies, we established the motile spontaneous mutant strain designated E22m1’. Filaments of the E22m1’ strain rapidly spread over the plate, and form a complicated morphology with mixture of dot and bundle of bacterial filaments (**Fig. 1A** middle left, **Movie S1**, upper right panel). Whole-genome sequencing identified five mutations on the genome in E22m1’ in addition to the kanamycin-resistance gene insertion (**Table 3**). A mutation was mapped to *LBDG_02920* encoding a putative DGC with insertion of IS200-like transposon (**Fig. 1B**). Another mutation was a silent mutation mapped onto *LBDG_25700* encoding an unknown function protein without changing amino acid sequences. The other three mutations were mapped to intergenic regions. Thus, we focused on the first mutation, namely, that on *LBDG_02920* and tested if this gene is responsible for the motility and morphological phenotype in E22m1’. In the *L. boryana* genome, we found nine genes encoding proteins harboring the GGDEF motif (**Fig. S1**). The nine genes were named *dgc1* to *dgc9* in order of the annotated nucleotide number on the genomic DNA, thereby designating the *LBDG_02920* gene, *dgc2*.

**Figure 1.**
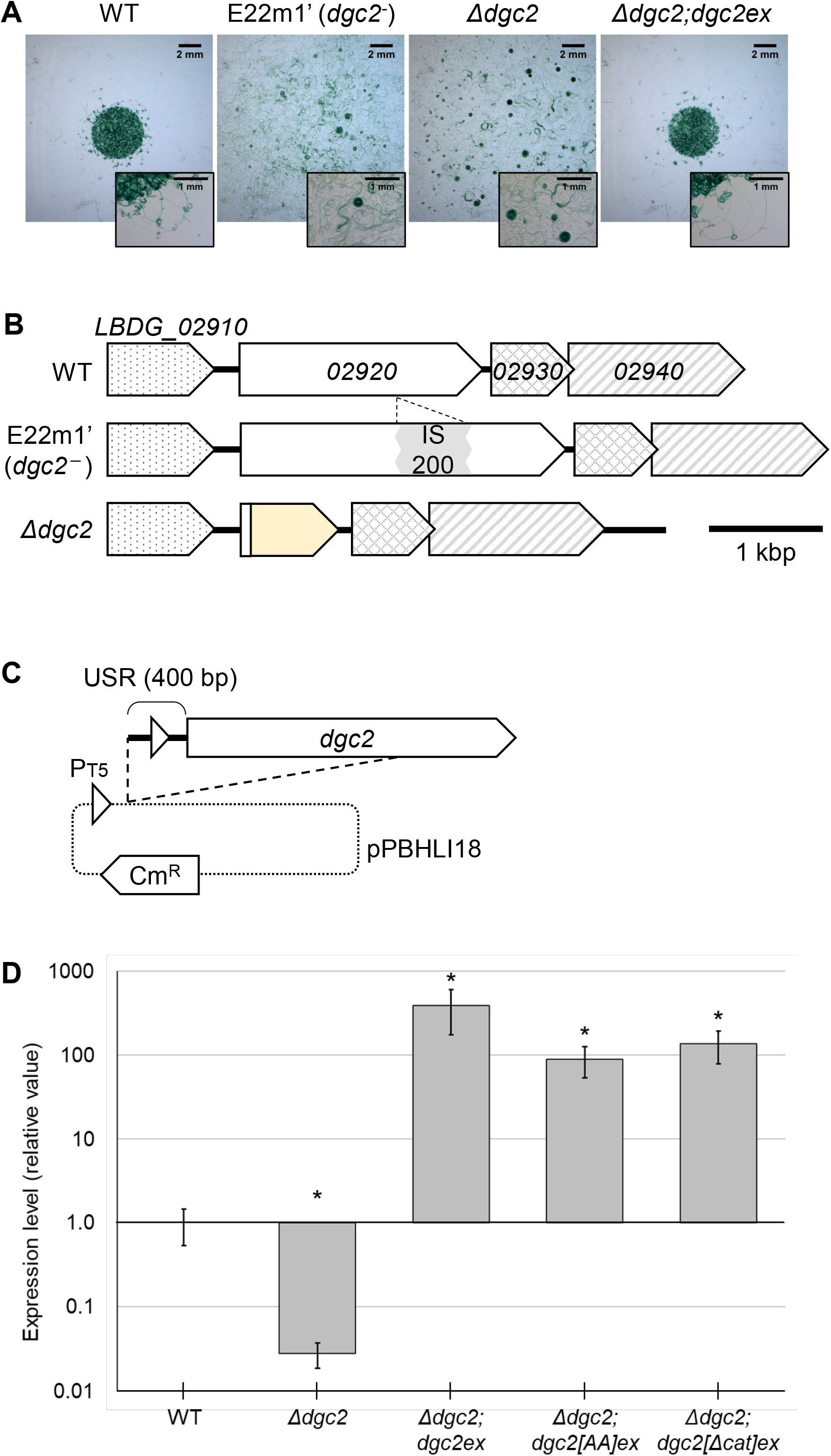
Disruption of *dgc2* allowed gliding motility and dot-like colony pattern formation in *L. boryana*. **A**. Morphological patterns of the wild type (WT; left), *dgc2*^*−*^(middle left), *Δdgc2* (middle right), and *Δdgc2; dgc2ex* strains on solid media at 9 days after inoculation of 3 μL each of cell suspension at the center of culture plates. Images with magnified scale are also shown at lower right of each panel. Both *dgc2*^*−*^ and *Δdgc2* strains showed gliding motility and spread rapidly on the solid media, and formed wandering and rotating clusters. **B**. Schematic representation of genomic DNA around the *dgc2* (*LBDG_02920*) locus of the wild type, *dgc2, dgc2*^*−*^ and *Δdgc2* strains. IS200-insertion in the *dgc2* open reading frame (ORF) in E22m1’ is shown in the gray box. Km^R^ (the yellow box) represents the kanamycin resistance gene. **C**. Schematic representation of the plasmid pIL1007 for ectopic *dgc2* expression. Triangles represent the T5 promoter (P_T5_) and a possible *dgc2* promoter within the 400-bp upstream region of the gene. Cm^R^ represents the chloramphenicol resistance gene. For details, see Materials and Methods. **D**. Expression profiles of the *dgc2* gene in strains by the qPCR analysis in the wild type, *Δdgc2*, and ectopically *dgc2*-derivative-induced strains. Asterisk represents a significant difference (*p*<0.05 Welch’s Wilcoxon rank sum test).

To validate if mutation in *dgc2* is responsible for motility, we substituted the *dgc2* open reading frame (ORF) with a kanamycin resistance gene (Km^R^) on the genome in the wild type (*dg5*) strain, resulting the *Δdgc2* strain (**Fig. 1B**). The *Δdgc2* strain regained gliding motility and morphological patterns essentially similar to that of E22m1’ on solid media (**Fig. 1A** middle right and **Movie S1** lower left panel). Then, we tried ectopic expression of *dgc2* (*dgc2ex*) in the *Δdgc2* background (*Δdgc2*;*dgc2ex*) to test if it complemented the phenotype in E22m1 (**Fig. S2A**). For overexpression, we modified a shuttle vector pPBHLI18 (39), harboring the replication origin of the pGL3 plasmid in *L. boryana* (40) and the *T5* promoter, which has been known functional for overexpression (**Fig. 1C**). As expected, *Δdgc2*;*dgc2ex* strain failed to show any signs of gliding motility, and the colony morphology bears a close resemblance to that of the wild type strain: cells grew staying at the inoculation point and showed ivy-like pattern at the edge (**Fig. 1A**, right panel, **Movies S1** lower right panel). The RT-qPCR analysis revealed that the expression level of *dgc2* with the ectopically expressed strains was ∼700-fold higher than that of the wild type strain (**Fig. 1D**). All these results are consistent with that motility and the colony pattern in E22m1’ is essentially due to nullification of *dgc2*; in other words, *dgc2* represses motility and formation of wandering clusters in *L. boryana*. Thus, hereafter the original mutant strain E22m1’ is referred to as the *dgc2*^*-*^ strain, which is different from the *Δdgc2* strain.

### Effect of *dgc2* on collective behavior

**Fig. 2** summarizes the collective behavior in the *Δdgc2* strain. Nine to twelve days after inoculation, bacterial filaments spread over tens of millimeters and showed a colony pattern with a characteristic cell distribution that was spatially biased with respect to cell density (**Figs. 2A** and **2B, Movie S2** left panel). Observed migrating high-density clusters are categorized into two groups: one is a ‘comet’-like cluster (comet, hereafter), which travels over solid media (**Fig. 3A, Movie S3**, left panel), while the other is a rotating disk-like cluster (**Fig. 3B, Movie S3**, right panel; hereafter disk). The disk kept moving to follow closed circular orbits. A kymograph shown in **Fig. 2B** indicates a time-dependent profile of the collective behavior and its transition. Vertical gradient lines on the kymograph (dark red and magenta arrowheads) represent rotating clusters that remain at the position. Many of the disks rotated steadily over 400 min as indicated by periodically fluctuating profile on the kymograph as is observed for *Pseudanabaena* sp. NIES-4403 (18) On the other hand, disk-like clusters in *Paenibacillus* sp. NAIST15-1 immediately stops rotating after forming large vortices (22). Collective behaviors in *Δdgc2* strain shown in **Fig. 2** are essentially similar to that in the original *dgc2*^*-*^ strain as shown in **Fig. S2** (**Movie S2**, right panel). These results suggest that *dgc2* in *L. boryana* is involved in suppression of the motility, thereby inhibiting the formation of migrating clusters.

**Figure 2.**
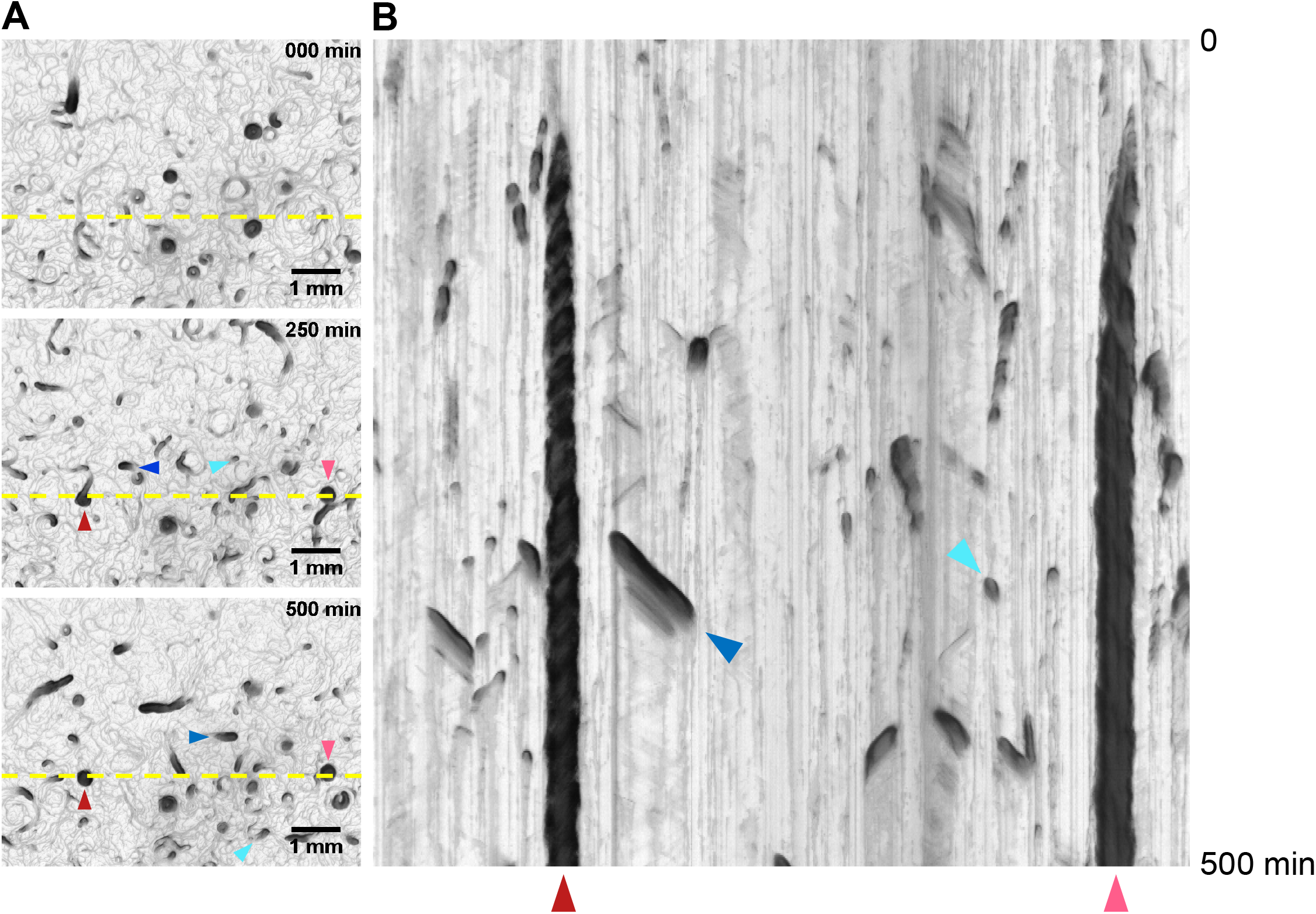
Transition of collective behavior of the Δ*dgc2* strain with time. **A**. Time-lapse images of *Δdgc2* strain on solid media. Time-lapse imaging was performed every minute, and the images were compiled into **Movie S2** (left). Time 0 (top panel) indicates the time that time-lapse imaging started, corresponding to ∼9 days after inoculation. **B**. A kymograph of colonies represented by yellow dashed lines shown in panel **A** over a 500-min period (top, min 0; bottom, min 500). Blue and cyan arrowheads represent comet-like wandering clusters, while red and magenta arrowheads represent disks. The comet represented by blue triangle appeared at the bottom of **Movie S2** (left) at 60 min, and glided near the yellow line from the 300 to 360 min, moving to the upper right. The comet indicated by the cyan arrowhead appeared at around hour 4 and passed through the line to the lower right on **Movie S2** (left). The disks appeared at around hour 2 and continued to rotate on the spot until the end of the imaging. In the last panel, designating 500 min, comets and two disks were also seeable.

**Figure 3.**
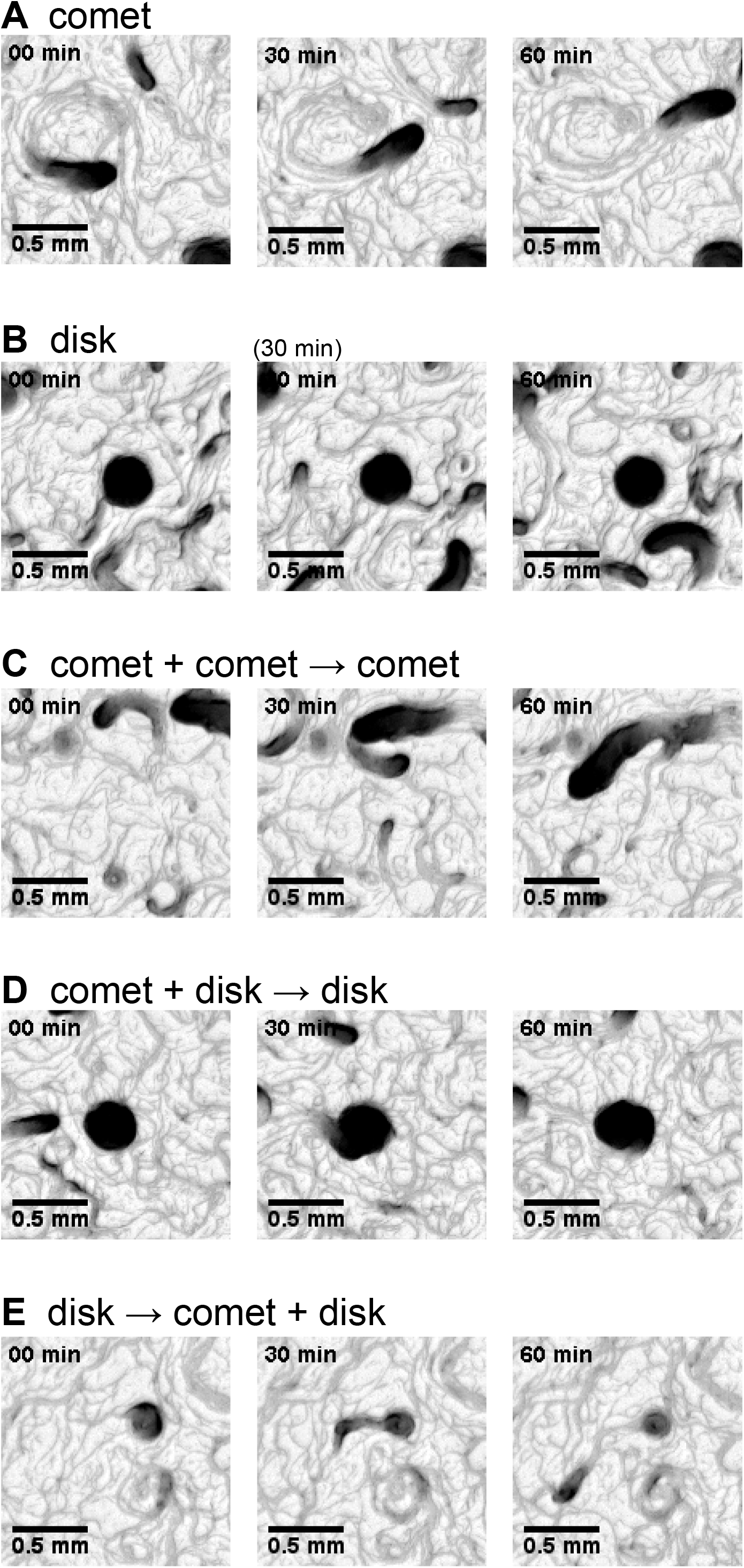
Time-course profiles of the wandering comet-like and rotating disk-like clusters. **A** and **B**. Movement and morphology of a comet (**A**) and a disk (**B**). **C**. Enlargement of a comet by unification of multiple comets. **D**. Enlargement of a disk by integrating a comet. **E**. Dissociation of comets from a disk.

Transiently emerging pattern of dark sloped short bars (cyan arrowhead) on the kymograph (**Fig. 2C** and **Fig. S3C**) indicate the passage of wandering comets. Although comet-like clusters in *Pseudanabaena* was proposed to be a pioneer in traveling to virgin area on the solid surface (18), in *L. boryana* comet-like cluster was observed to move exclusively on already existing passages. Their speed of movement is not constant, and some clusters were observed to collapse suddenly. Thus, it seems likely that comets are more fragile and more easily affected by microenvironmental conditions in *L. boryana*, compared with that in *Pseudanabaena*.

We have observed some transition of clusters with collision. **Fig. 3C** and **Movie S4A** show unification of multiple comets. In this case, a preceding small comet had initially formed a circular orbit. When another large comet entered from behind, crossing the orbit. The small comet kept moving and then caught up with the slowly moving large comet to be unified. In the case of collision of a comet and a disk shown in **Fig. 3D** and **Movie S4B**, a comet moved to the direction of the tangent line of the disk’s rotation, and was smoothly unified into the rotating disk. We also observed a disk-to-comet transition. As shown in **Fig. 3E** and **Movie S4C**, a group of comets sneaked out from inside of the disk and moved straight, while they formed new small closed orbit within 1 h. The original disk continued to rotate slowly until the end of the observation. After the divergence of the comet-like clusters, the uniformity of the rotational direction was broken, and the migrating direction of the subpopulation in the disk was at least transiently disordered. This is probably because impact of the influx and outflow of the filaments into/from the disk became relatively higher, and the uniformity of the migrating direction was less maintained.

### Counterclockwise movement is more stable for high density clusters

For more detailed observations, we analyzed behavior of individual clusters shown in **Fig. 2** and **Movie S2**. Interestingly, the rotation direction of the stably rotating disks was mostly counterclockwise (CCW), for observers looking at the agar surface from the top, as exemplified by a disk shown in **Fig. 4A** and **Movie S5A**. Since the cell population inside the disk is not uniform, the density on the kymograph (**Fig. 4, Fig. S3**) changes little by little as the disk rotates; if it rotates in the CCW direction, it appears as oblique lines to the left on the upper side of the disk (yellow) and to the right on the lower side of the disk (blue). It is the case for the two disks on the kymograph shown in **Fig. 2B**, where the reference line (yellow line on **Fig. 2A**) crossed the top of the left disk and the bottom of the right disk, thereby generating oblique lines to the left and right, respectively. The disk in **Fig. 3B, 3D**, and **4A**, and **Movie S5A** kept rotating stably in the CCW direction for 300 min, even though the disk collided with comet several times, which corresponds to the temporary increase in the width of the black line on the lower (blue) kymograph (**Fig. 4A**). On the other hand, another disk shown in **Fig. S3A** and **Movie S5B** rapidly disintegrated after ∼ 150-min CW rotation with separation of a small comet-like cluster from the edge of the disk. A disk shown in **Fig. 4B** and **Movie S5C** was rotating stably CCW about 350 minutes (indicated by the red arrowhead), but the direction of rotation changed to CW. Then, within 150 min the disk began to disintegrate with separation of the limb population at the edge of the disk by forming a comet-like small cluster. However, a change in the direction of rotation to CW does not necessarily mean that the disk will change into a comet. For example, a disk shown in **Fig. S3B** and **Movie S5D** initially rotated CCW, but after 840 minutes (green arrowhead), most of the cells from the edge were separated from the disk in the form of comet, and the remaining center maintained its rotation for a while, but stopped after about 30 min. Such a case of disintegration while maintaining CCW rotation is exceptional. In more details, at the initial phase (0-90 min in **Fig. S3B** and **Movie S5D**) many filaments were passing through a hollow circular orbit, and then the diameter of the hollow suddenly shrank at around 90-120 min to form a rotating disk. Other comets collided with the disk at around 185 (red arrowhead) and 400 min (blue arrowhead), and a group of comets popped out at around 300 min (yellow arrowhead). It looks likely that the impact of the collision at ∼400 min was so significant that the shape of the disk became a distorted oval. After 840 min (green arrowhead), a disk ejected a comet that split itself into two parts, as shown in **Fig. 3E** and **Fig. S3B**. This result seems to suggest that even if the rotation is in the CCW direction, when too large a perturbation is given and the angular velocity of the disk is extremely unbalanced, the disk may collapse.

**Figure 4.**
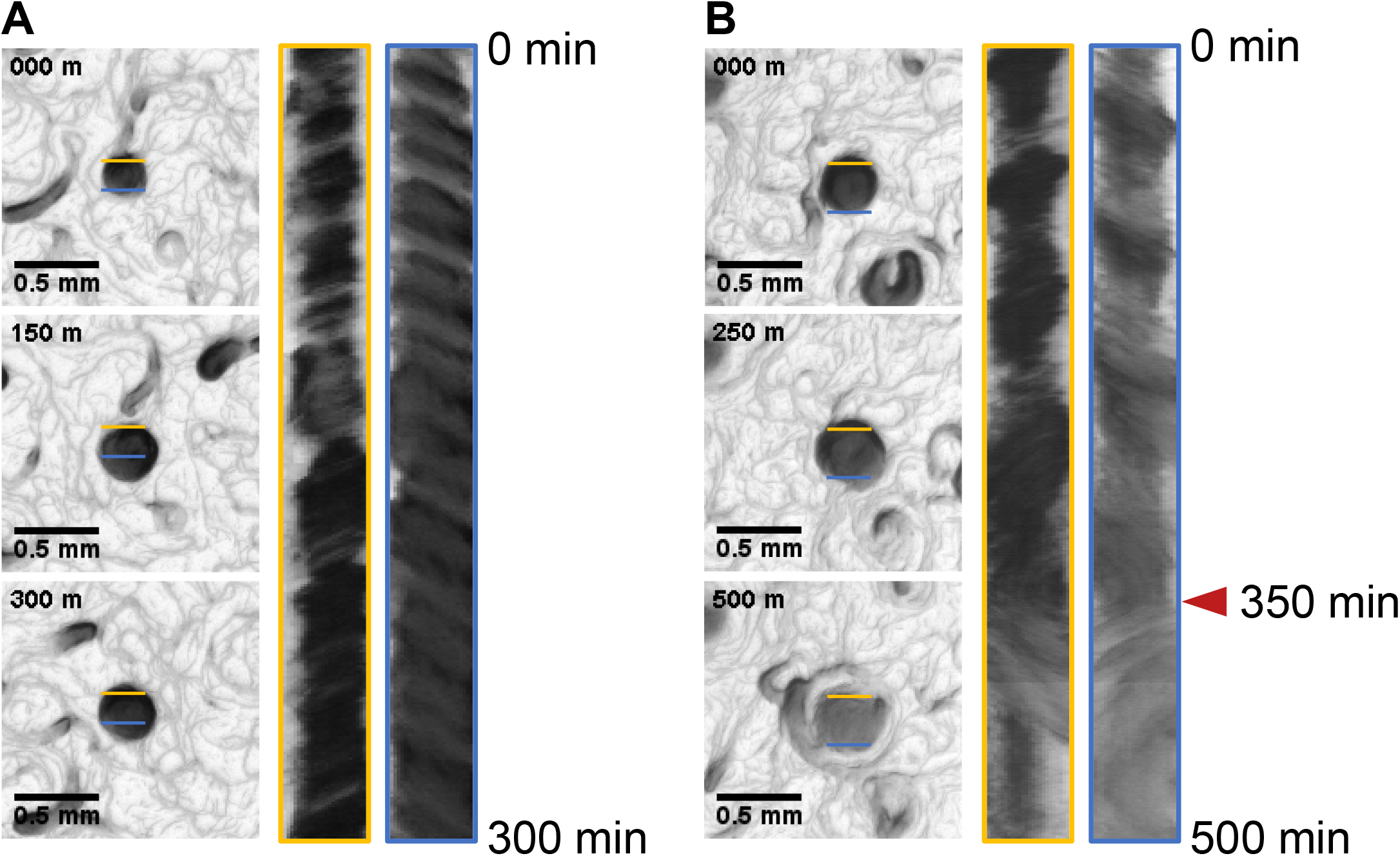
Counterclockwise preference of the disc rotating stably. **A**. kymograph representing stable rotation of a disk for more than 300 min, while this disk collided with three comets at around min 0,150, and 300. The left (yellow) and right (blue) kymographs represent time-course images at the upper side and the lower side of the disk indicated with the corresponding color, respectively. **B**. Disintegration of a disk, accompanying by a CCW-to-CW shift in the direction of rotation.

Considering the CCW-preference in disk-like clusters, we analyzed whether comet-like clusters also show a CCW or CW bias in the direction of motion. For time-lapse images of comets, the angle (*θ*_*t*_) of the velocity vector was obtained from the X-axis direction, and the angle difference (*Φ*_*t*_) between the angle (*θ*_*t*_) and next angle (*θ*_*t+1*_) was calculated. This *Φ*_*t*_ values were used as indices of the perturbation angle (**Fig. 5A**). By calculating the duration for which the *Ft* values of taking the same sign (+ or -), we estimated how long the directionality of the motion is maintained. For example, **Fig. 5B** indicates trajectory of the migration of a comet within 663 min, and **Fig. 5C** represents the distribution of the duration of moving in the same direction and the average *Φ*_*t*_ values during that time. **Fig. 5D** summarized the results obtained from 19 independent comets. Apparently, long-lasting (8-to-12 min) directional movement was exceptionally limited to CCW, while the motion in the CW direction lasted for a maximum of only 7 min. Although the distribution of probability density for median perturbation angle and duration showed no significant difference in the distribution of median perturbation angle between CCW and CW (Mardia-Watson-Wheeler test, *p* = 0.6369), the duration of the CCW direction was statistically longer than that of the CW direction (Student’s *t*-test, p << 0.01). Interestingly, *Flavobacterium johnsoniae* was also recently shown to perform gliding motility and vortex formation with the CCW preference (6). It shows a CCW trajectory even from the single-cell state, and remains unified in the CCW direction while rotating as a group, but moves in the CW direction just before it stops its rotation. n addition, asymmetric movement for asymmetric colony pattern formation has been found in *Paenibacillus*, which moves with a swarming motility rather than a gliding motility (41). The mechanism of the preference for CCW in *L. boryana* remains to be solved.

**Figure 5.**
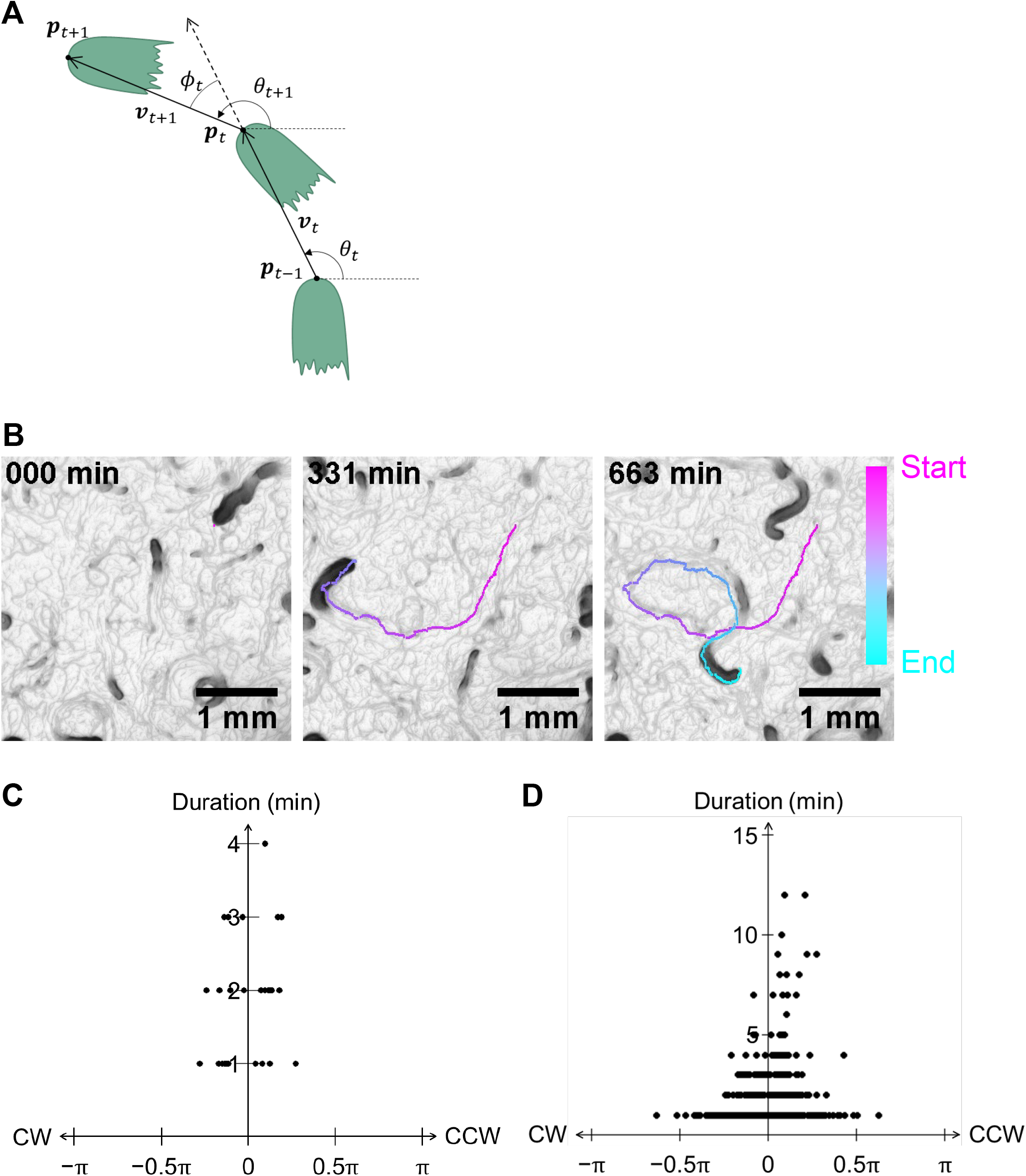
Counterclockwise preference of the comet. **A**. Schematic diagram of the angular velocity of a comet. *p_t_*, position of a head of a comet at time *t*; 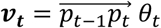; the angle between the velocity vector ***v***_***t***_ and the x-axis; *ϕ_t_* = *θ*_*t*+1_ − *θ*_*t*_, the angular velocity. **B**. Trajectory of a comet. Colors indicate the positions of the cluster at min 0 (magenta) and 663 (cyan). **C** and **D**. The distribution of the duration of movement in the same direction and the average *ϕ*_*t*_ values in the comet indicated in panel **B (C)** or that in 19 independent comets (**D**).

### Extracellular polysaccharides secretion

Pilli and the secretion of extracellular polysaccharides (EPS) are known to be involved in cyanobacterial gliding activity (8, 9, 17, 189). In *Oscillatoria* and *N. punctiforme*, especially, staining of the trajectory of the filaments as they pass through with india ink has been established to show remaining mucus-like substance on trails (9, 42; 43;). The presence of EPS was confirmed in *N. punctiforme* and *Pseudanabaena* using fluorescent dye-conjugated lectin subunits, RCA-120 (9, 18). Thus, we examined both the india ink assay and the RCA-120 staining in *L. boryana*. We cultured the *Δdgc2* strain on BG-11 solid media for several days and allowed it to glide on the surface. They were then stained with india ink and observed under the microscope for 12 min (see Materials and Methods). As shown in **Fig. 6A** and **Movie S6**, the trails were stained with india ink particles, confirming the gliding filaments secreted viscous materials. Next, we tested if the *Δdgc2* population was stained with RCA-120-conjugated to fluorescein that binds to galactose and *N*-acetyl galactosamine residues. Positive RCA-120 signals were visualized around the filaments, confirming the EPS secretion in the *Δdgc2* strain (**Fig. 6B**). On the contrary, positive signals were hardly observed in the non-motile wild-type strain (**Fig. 6B**). These results strongly suggest that the gliding activity in *L. boryana* is activated by EPS-based mucus, and DGC2-derived c-di-GMP production prevents the EPS secretion.

**Figure 6.**
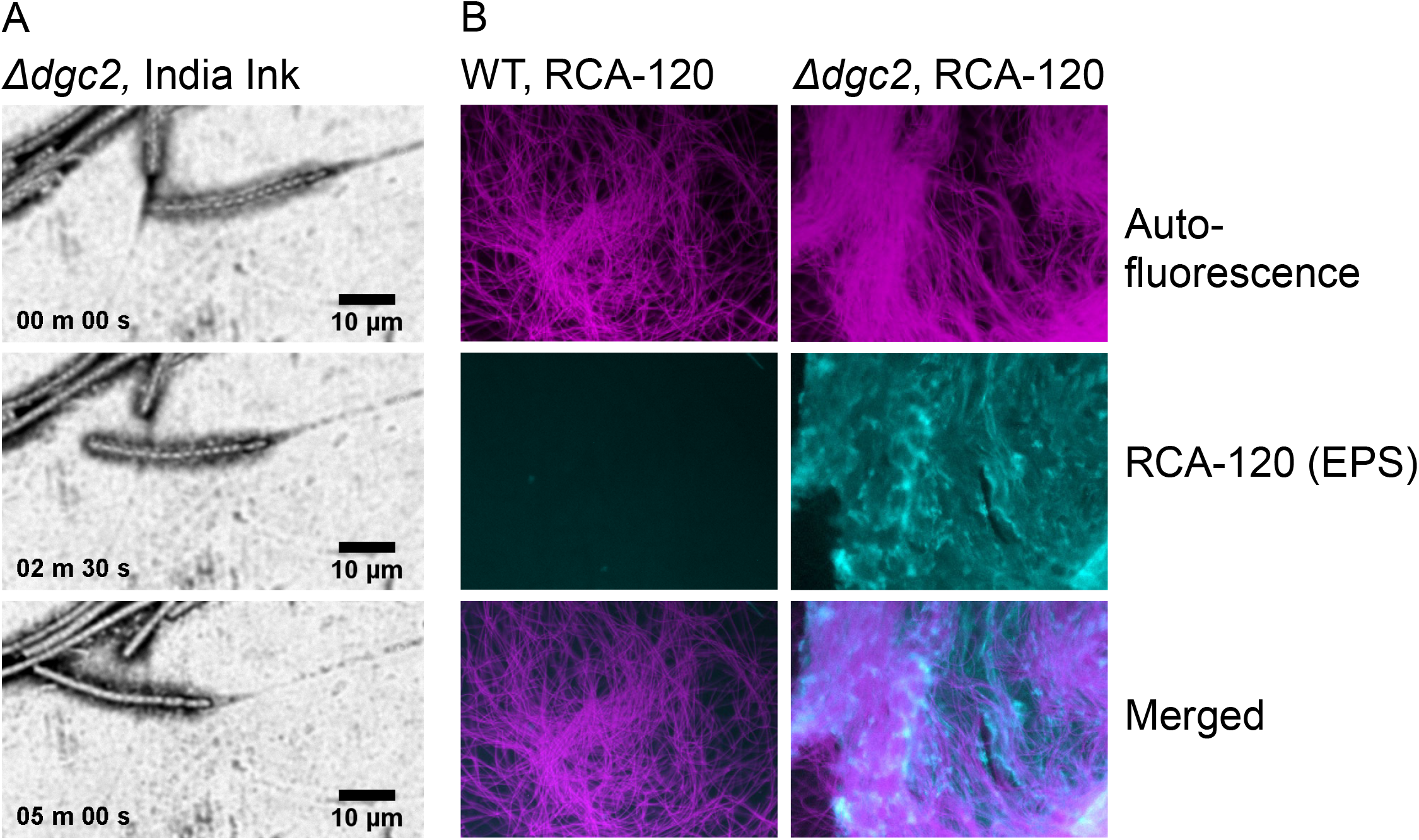
The *Δdgc2* strains secretes EPS. **A**. India ink-staining of viscous trails of the *Δdgc2* filaments. The video is provided as **Movie S6. B**. Fluorescent microscopic images of RCA-120 stained *Δdgc2* cells. Magenta and cyan indicate autofluorescence of cells and stained EPS, respectively.

### GGDEF domain of Dgc2 is important for inhibiting motility and high-density cluster formation

Dgc2 is consist of CHASE2, transmembrane motifs, PAS, PAC, and the GGDEF domain from the N-terminus (**Fig. 7A**, top). CHASE2 (cyclase/histidine kinase-associated sensing extracellular-2) is an extracellular module conserved among bacterial sensory proteins, while its environmental ligands remain unknown (44. 2003). The PAS (Per-Arnt-Sim) domain is a module for protein-protein interaction or binding to ligands, while PAC (PAS-associated, C-terminal) motif contributes to folding of the PAS domain (45). As mentioned above, overexpression of *dgc2* (*dgc2ex*) suppressed the motile activity of *Δdgc2* (**Fig. 1D**). In order to validate the significance of the GGDEF domain, we tested overexpression of two mutant derivatives of *dgc2* to see if it failed to inhibit motility. As represented in **Fig. 7A** and **7B**, one derivative is an active-site mutant in which GGDEF was substituted with GGAAF (Dgc2[AA]) and the other lacks the GGDEF domain (Dgc2[Δcat]). As expected, both of the resulting mutant-overexpressor strains, *dgc2[AA]ex* and *dgc2[Δcat]ex*, failed to inhibit motility in *Δdgc2* background (**Fig. 7C**). However, their phenotype was not completely same to the *Δdgc2* strain. Compared with the *Δdgc2* strain, the mutant-overexpressor strains tended to remain at the initial position of inoculation, while outward expansion of the colony with nematic alignment was evident (**Fig. 7C**). Thus, the phenotype of these mutant-overexpressor strains is likely intermediate between the wild type and the *Δdgc2* strain. Consistently, under this experimental condition with illumination of 7.5 μmol m^-2^ s^-1^, overexpression of mutated Dgc2 failed to develop high density cluster formations, different from the *Δdgc2* strain (**Fig. 7C**). On the contrary, under a bit higher light intensity (10 μmol m^-2^ s^-1^) overexpression of either *dgc2[AA]* (**Fig. 7C**) led to comet and disk formation, while that of the intact *dgc2* strikingly suppressed motility (**Fig. 7C**). Although the reason of this light intensity dependency remains unknown, all these results are consistent with functional importance of the GGDEF motif or DGC activity in DGC2 for inhibiting motility and migrating cluster formation. Some difference between the mutant overexpressor and *Δdgc2* strains could be due to dominant-negative effect of the overproduced N-terminal portion of DGC2 on other CHASE2- or PAS-related proteins.

**Figure 7.**
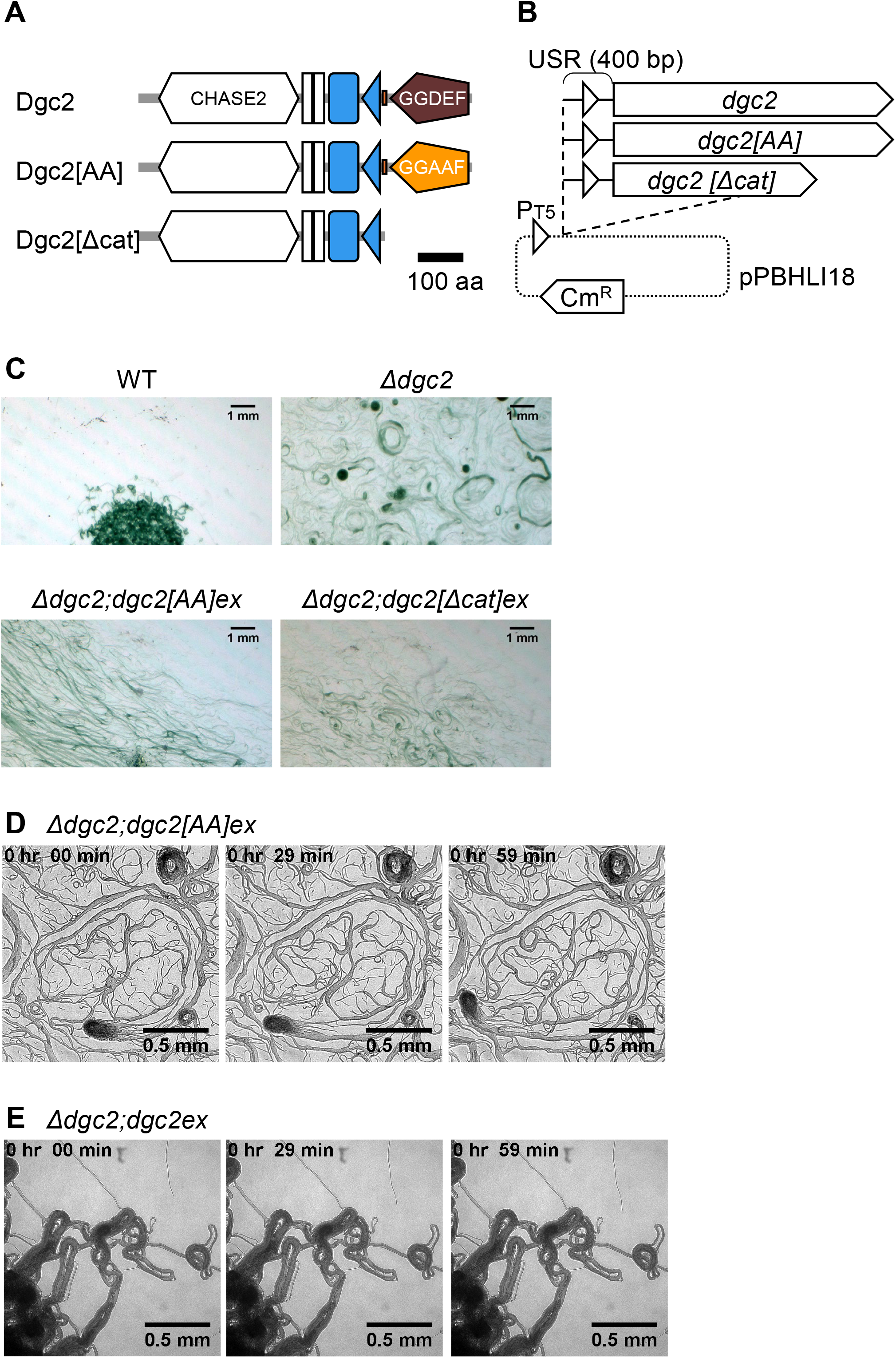
GGDEF domain of Dgc2 is important for inhibiting motility and pattern formation. **A**. Schematic representation of the wild type and mutant Dgc2 variants for ectopic expression. CHASE2 domain (white hexagon), transmembrane motifs (white squares), PAS domain (blue square), PAC domain (blue triangle), and the GGDEF motif (brown pentagon) are shown. **B**. Schematic representation of plasmids for overexpressing *dgc2[AA]* and *dgc2[Δcat]* as represented in **Fig. 1C. C**. Morphological colony patterns in the mutant overexpressor strains. Each cell suspension was spotted onto solid media at similar position (middle low of each panel) with illumination of 7.5 μmol m^-2^ s^-1^, and images are captured x days after inoculation. **D** and **E**. Movement of the *Δdgc2-*background strains which overexpress either *dgc2*[*AA*] (**D**) or intact *dgc2* (**E**) with illumination of 10 μmol m^-2^ s^-1^ for 1 h.

### Dgc2 inhibits biofilm formation in *L. boryana*

c-di-GMP is known to facilitate biofilm formation in many species of bacteria (35). Thus, we examined if biofilm formation is affected in the presence or absence of *dgc2* in *L. boryana*. Initially, *L. boryana* was grown in BG-11 liquid media on flat bottom glass flasks (diameter of 60 mm) with or without aeration under different light conditions. We found that the wild type cells formed some aggregates, and cells tended to sank to the bottom reproducibly under low light conditions of less than 10 μmol-m^-2^-s^-1^ with gentle aeration of the air (**Fig. 8A**, WT with “−”). However, when liquid media were sucked out using a pipette, cells rarely retained, indicating that cells did not tightly adhere to the bottom (**Fig. 8A**, WT with “+”). Some filaments also formed some aggregates at the air-liquid interface (flocculation), while the aggregates were very fragile to be easily disrupted even by gentle holding up the flask. The aggregates (or flocculates) formed inside the flask were also fragile so that they were easily broken during pipetting or decantation of the medium. On the other hand, the *Δdgc2* strain developed more larger aggregates at the bottom (**Fig. 8A**, *Δdgc2* with “−”). The aggregates were so firmly attached to the bottom as stable biofilm that it was difficult to disappear by sucking the liquid (**Fig. 8A**, *Δdgc2* with “+”), even after vigorously rubbed with the tip of a pipette. It was surprising because nullification of DGC was expected to reduce c-di-GMP content, thereby inhibiting biofilm formation. We next quantified the ratio of bottom-attaching biofilm mass to total cell mass (adhesion rate), employing the same method used for *Synechococcus* biofilm formation (46). Cells were cultured in 10 ml of BG-11 medium on a glass-bottom plates with a dimeter of 60 mm to facilitate biofilm formation for 10 days. Then, the supernatant was collected by pipetting and subjected to chlorophyll extraction with methanol. The resulting value of chlorophyll content was represented as *Cs*. Chlorophyll was also extracted from tightly glass-attached cells (biofilm), and its content was also quantified as *Cb* (**Fig. 8B**). Despite a large variation in the total chlorophyll content (*Cs + Cb*) among the plates, the general trend was that the *Cb* value in the *Δdgc2* strain was higher than that of the wild type strain (**Fig.8C, Fig. S4**). Particularly, it seems apparent that the difference increases with increasing *Cs* value, strongly supporting a density-dependent effect in which biofilm formation on the glass bottom is promoted as cell density increases. Under the microscope, *Δdgc2* bacterial filaments were scattered on the glass surface, forming foci (aggregates) of various sizes (**Fig. 8D**). The foci were formed with many filaments or bundles, extending radially outward according to their size, and the foci were connected to each other through the filaments. This morphology resembles the structure of a scale-free network of interconnected hubs of various sizes, suggesting that cell aggregation during biofilm formation involves multiplication processes characteristic of such networks. On the other hand, the wild type filaments rarely formed such structure, while we still observed a faint bundle or single filaments attached onto the surface more randomly (**Fig. 8D**).

**Figure 8.**
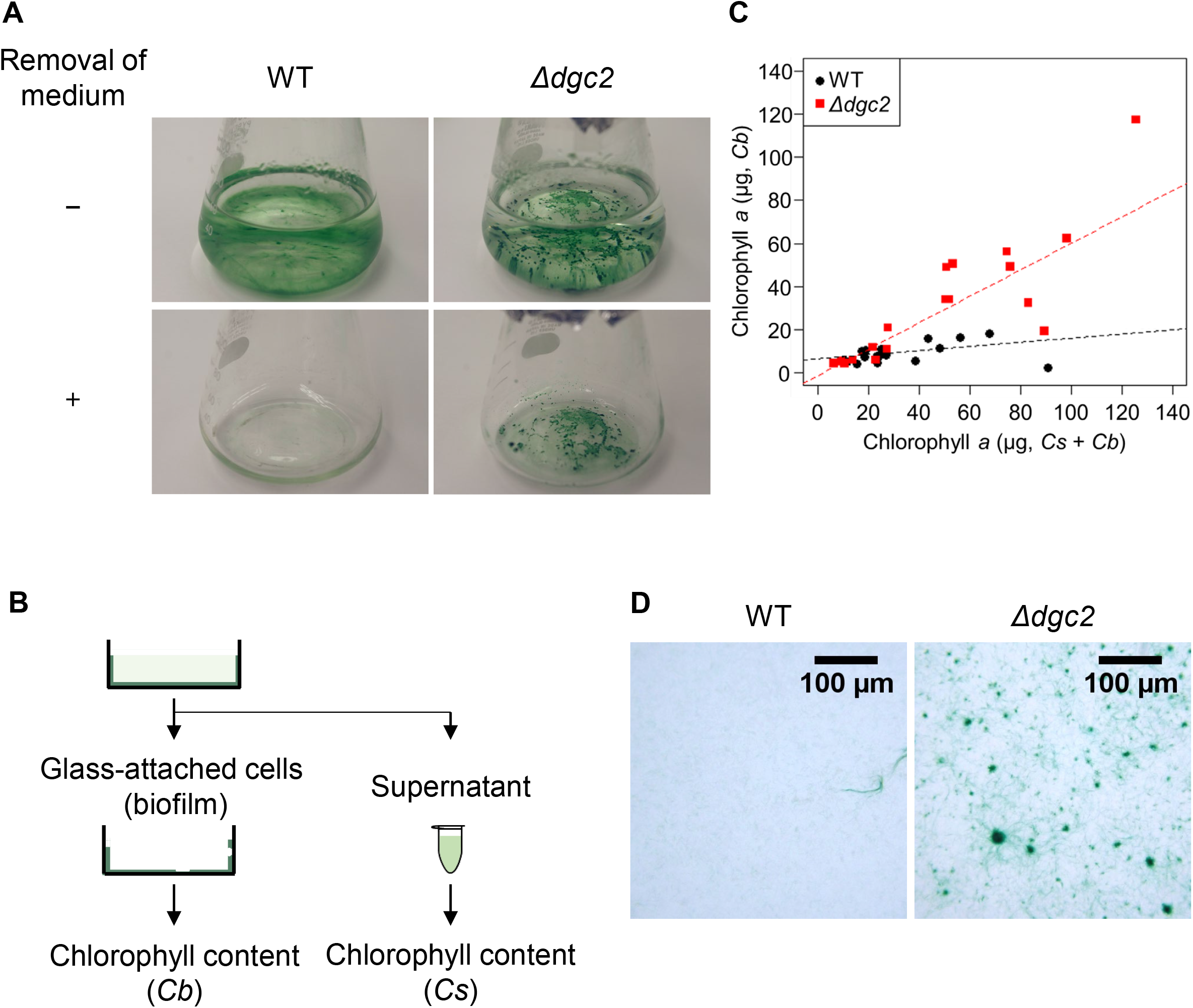
Dgc2 inhibits biofilm formation in *L. boryana*. **A**. Representative images of aggregation/biofilm profiles in the wild-type and *Δdgc2* strains before (-) and after (+) removal of liquid media. The wild type formed some floating aggregates which were not tightly attached to the glass surface. On the other hand, the *Δdgc2* strain developed larger aggregates which attached more tightly to the bottom (biofilm). **B**. Schematic representation of the experiment to quantify the ratio of bottom-attached cells (biofilm) to the total cell mass (adhesion rate, *Ra*). Cells were cultured in 10 mL of BG-11 medium in a glass-bottom plate. Then, supernatant was transferred to another tube gently. Chlorophyll content in the supernatant and biofilm were represented as *Cs* and *Cb*, respectively. **C**. Distribution of the total chlorophyll content (*Cs* + *Cb*, abscissa) and the chlorophyll content extracted from glass-attached cells (*Cb*, ordinate) in each experiment. Black and red plots indicate data for the wild type and *Δdgc2* strains, respectively (*n*=20 for both strains). Dotted lines indicate linear approximate lines with correlation coefficients (Spearman’s rank correlation coefficients) of 0.76 and 0.55 for the *Δdgc2* and the wild type strains, respectively. **D**. Microscopic images of glass-attached cells in the wild type and *Δdgc2* strains, after discarding the liquid phase.

## Conclusion

We have identified that Dgc2 suppresses motility in *L. boryana*. Although motility has not been reported in the wild type *L. boryana* strain, some species belonging to *Leptolyngbya* are known to show gliding activities (47). Our results do not exclude a possibility that the wild type *L. boryana* may have a potential to show motility if c-di-GMP production by Dgc2 is lowered under as-yet-unknown natural condition. The presence of the CHASE2 domain and PAS domain in Dgc2 (**Fig. 7A**) would support this possibility: sensing an extracellular environmental signal at CHASE2 and/or binding a cognate partner at PAS domain would control the enzymatic activity. Alternatively, the wild type strain which may have already obtained additional spontaneous mutations to suppress motility during maintenance at laboratories for many decades. In any cases, searching for possible ligands or binding partners of Dgc2 should be followed to address this question. It should be noted that genes encoding Dgc2-like, CHASE2-containing DGCs are found not only in cyanobacteria but also widely in genomic DNAs of other bacterial species belonging to Proteobacteria (such as *Desulfobulbaceae*) and Verrucomicrobia (such as *Methylacidiphilum*) (**Table S3**). Thus, CHASE2-mediated regulation of c-di-GMP signaling seems widely conserved. However, it is not entirely clear what ligands and signals are accepted by CHASE2 in these DGCs. The genetically tractable *L. boryana* should provide an excellent model to address this issue.

We also revealed that *L. boryana* Dgc2 suppressed biofilm formation (**Fig. 8**), in contrast to some previous reports that c-di-GMP or DGCs facilitate(s) biofilm formation in many bacterial species, represented by *Pseudomonas aeruginosa* (47). This interesting difference suggests the presence of a variety of relationships between biofilm formation and motility through c-di-GMP regulatory network. For example, in *Pseudomonas* motility may act as an inhibitor of biofilm formation by decreasing the strength of cell-cell interaction through the movement of individual cells. On the other hand, in the *Δdgc2* strain of *L. boryana*, the acquisition of motility may rather promote migration-associated grouping and thus enhance the strength of the interaction among filaments. If the latter is true, then formation of the high-density migrating clusters observed on the agar medium could be interpreted as a pattern formation phenomenon brought about by the physical properties of biofilm formation with motility under the artificial conditions of the flat solid surface of agar media. The formation of foci by mobile filaments for biofilm formation shown in **Fig. 8D** also supports this possibility.

## Material and Methods

### Strains and culture conditions

Strains used in this study were summarized in **Table 1**. *Leptolyngbya boryana dg5* strain which harbors heterotrophic capability under continuous dark conditions (Fujita *et al*., 1996, 27) was used as a wild type strain. The original motility strain E22m1’ was isolated as a spontaneous mutant, derived from a transformant strain, E22m1-dg5. The E22m1-dg5 parental strain carried a kanamycin resistance gene (27-bp downstream of the *LBDG_21990* ORF) which did not show any observable phenotypes including motility nor colony morphology. The motile E22m1 strain was renamed *dgc2*^*-*^ strain after genetic analysis described herein. For generating *dgc2-*depleted strains, we connected a1500-bp upstream segment of *LBDG_02920*, a kanamycin-resistance gene cassette from pYFC10 (26), and a 1500-bp downstream segment of *LBDG_02920* fragments in this order and then cloned into a 1752-bp segment derived from pBluescript II SK(+) (Stratagene) to give rise to pIL910. For *dgc2*-overexpressor strains, we cloned a segment covering a 400-bp upstream region of *LBDG_02920* and the coding open reading frame region into the *Sph*I-*Bam*HI site of a shuttle vector, pPBHLI18 (39) to generate the resulting plasmid pIL1007. pPBHLI18 is a derivative of pPBH201 (40), harboring the *T5* promoter and terminator derived from pQE70 (Qiagen), the replication origin of the plasmid pGL3 (Promega) in *L. boryana*, and a chloramphenicol-resistance gene cassette. Suttle vectors harboring mutated *dgc2* gene (pIL1018 and pIL1019) was generated by PCR-based targeted mutagenesis of pIL1007. DNA fragments for these constructions were generated by PCR with primers listed on **Table S1**. Connecting the DNA segments was performed using the NEBuilder HiFi DNA Assembly Kit (NEB) or Ligation high Ver.2 (TOYOBO). The list of resulting plasmids was summarized in **Table 2**. All plasmids were checked by DNA sequencing after completion. Transformants used in this study was generated with the electroporation method (26, 27, 48) with some modification such that we centrifuged for collection and washing of cells, instead of the filtration method described by Tsujimoto *et al*. (48). Deletion of genes was confirmed by Southern blotting analysis using the DIG DNA Labeling Mix (Sigma-Aldrich). Transformation with shuttle vectors was confirmed by PCR using purified genomic DNA as each template. All strains were cultured in BG-11 solid or liquid media (49) supplemented with 7.5 mM glucose under illumination. We set white fluorescence lamps of 7.5 μmol m^-2^ s^-1^ for most experiments, except for r experiments shown in **Fig. 6, Fig. 7D** and **7E** and **Movie S6** at 10 μmol m^-2^ s^-1^ and experiments shown in **Fig. 6** and **Fig. S3** at 15 ∼ 20 μmol m^-2^ s^-1^. 20 μg/mL of kanamycin or 10 μg/mL of chloramphenicol was added if necessary. To compare the colonies formed by each strain (**Figs. 1A, 2 to 4, 5A, 7C, 7D**, and **7E, Figs. S2** and **S3**, and **Movies S2** to **S5**), we spotted the bacterial suspensions on agar medium. Cells grown on agar plates were collected with a platinum loop, suspended in BG-11 liquid medium to OD_730_ of 0.1, and 2-μL cell suspension was inoculated on a fresh BG-11 agar plate for about 2 weeks under low light illumination of about 7.5 μmol m^-2^-s^-1^. The strains harboring the shuttle vector plasmid were cultured on BG-11 agar media containing chloramphenicol, while other strains were cultured on BG-11 agar medium without antibiotics. At the time of inoculation, the wild type and *Δdgc2* strains were always cultured as controls.

**Table 1.**
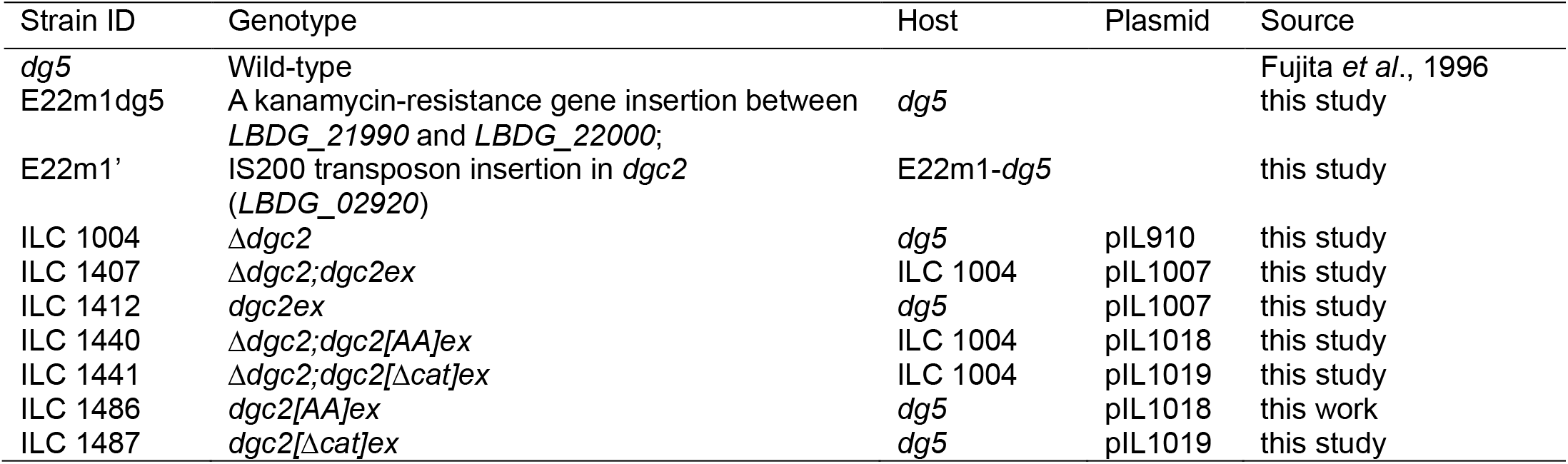
Strains used in this study.

| Strain ID | Genotype | Host | Plasmid | Source |
| --- | --- | --- | --- | --- |
| *dg5* | Wild-type |  |  | Fujita *et al.*, 1996 |
| E22m1*dg5* | A kanamycin-resistance gene insertion between *LBDG_21990* and *LBDG_22000*; | *dg5* |  | this study |
| E22m1' | IS200 transposon insertion in *dgc2* (*LBDG_02920*) | E22m1-*dg5* |  | this study |
| ILC 1004 | $\Delta dgc2$ | *dg5* | pIL910 | this study |
| ILC 1407 | $\Delta dgc2;dgc2ex$ | ILC 1004 | pIL1007 | this study |
| ILC 1412 | *dgc2ex* | *dg5* | pIL1007 | this study |
| ILC 1440 | $\Delta dgc2;dgc2[AA]ex$ | ILC 1004 | pIL1018 | this study |
| ILC 1441 | $\Delta dgc2;dgc2[\Delta cat]ex$ | ILC 1004 | pIL1019 | this study |
| ILC 1486 | *dgc2[AA]ex* | *dg5* | pIL1018 | this work |
| ILC 1487 | *dgc2[\Delta cat]ex* | *dg5* | pIL1019 | this study |

**Table 2.** Plasmids used in this study.

| Plasmid | Construction | Origin | Resistance | Host vector |
| --- | --- | --- | --- | --- |
| pIL910 | USR 1500 bp of *dgc2::KmR::DSR* 1500 bp of *dgc2* | ColE1 | Ap<sup>R</sup>, Km<sup>R</sup> | pBlueScript SK II (+) |
| pIL985 | *Sph* I-*dgc2-Bam*H I | ColE1 | Cm<sup>R</sup> | pPBHLI18 |
| pIL1007 | *Sph* I-USR400 bp of *dgc2::dgc2-Bam*H I | ColE1 | Cm<sup>R</sup> | pIL985 |
| pIL1018 | *Sph* I-USR400 bp of *dgc2::dgc2[AA]-Bam*H I | ColE1 | Cm<sup>R</sup> | pIL1007 |
| pIL1019 | *Sph* I-USR400 bp of *dgc2::dgc2[\Delta cat]-Bam*H I | ColE1 | Cm<sup>R</sup> | pIL1007 |

**Table 3.** Polymorphism between WT (*dg5*) and E22m1’ (*dgc2*^*−*^) genome.

| Nucleotide loci on the genome | ORF ID | Putative function | Mutation | Mutation on protein | Length of coding protein (residues) |
| --- | --- | --- | --- | --- | --- |
| 324676-324948 | *LBDG_02920* | Diguanylate cyclase | IS200 insertion | Leu456fs | 732 |
| 2754184 | *LBDG_25700* | Hypothetical protein | A to G substitution | Gly76Gly | 183 |
| 1535162-1535163 |  | Non-coding region | 2-base (AA) deletion |  |  |
| 2368755 |  |  | Neomycin phosphatase (Kanamycin resistance gene) |  |  |
| 2431019-2431020 |  | Non-coding region | 1-base (T) insertion |  |  |
| 2630555-2630556 |  | Non-coding region | 1-base (A) insertion |  |  |

Resequencing of the genomic DNA from E22m1' (*dgc2*<sup>-</sup>) revealed polymorphism at six loci including a frameshift mutation caused by the insertion of the IS200 transposon in *LBDG\_02920* (Leu456fs) and insertion of the neomycin phosphatase (kanamycin resistance) gene at genomic locus of 2368755. The insertion of the kanamycin resistance gene was artificially carried out during the isolation of the mutant E22m1, which was the source of the motility mutant strain. Another nucleotide substitution of adenine with guanine was found in the *LBDG\_25700* ORF at a genomic locus of 2754184, which is a silent mutation that did not change the glycine residue. Any of the other mutations were found in intergenic regions: deletions of two bases (adenines) at locus of 135162 and 153563, thymine-insertion between 2431019 and 2431020, and adenine-insertion between 2630555 and 2630556.

### EPS staining assay

EPS staining with fluorescent lectin (RCA-120) was performed as described previously (9, 50) with slight modification as follows (**Fig. 6B**). One hundred μL of cell suspension was centrifugated at 3,500 rpm, at room temperature (RT) for 5 min and the supernatant was removed to collect cell pellets. Then, 100 μL of fresh BG-11 medium containing RCA120-fluorescein (Vector laboratories, USA) at 20 μg/ml (final concentration) was added to the collected cells and incubated at RT for 30 min. After centrifugation at 3,500 rpm at RT for 5 min, the supernatant was removed., and the cell pellets were suspended in 100 μL of fresh medium and then placed on a slide glass. The stained materials were observed under the IX-71inverted microscope (Olympus, Japan) with LUCPlanFLN (20x) objective (Olympus, Japan) and filter sets for RCA120-fluorescein (Ex 485-517 nm, Em 505-565 nm) and for autofluorescence (Ex 544-592 nm, Em 570-650 nm). For ink staining, cells cultured on BG-11 solid media for about 1 week were used. 10 μL of solution containing 2% india ink (Winsor & Newton, UK), 4 μM CaCl_2_, and 0.05% (v/v) Triton X-100 (Sigma) was then dropped on the surface of the medium. Images were taken under microscope (IX71; Olympus, Japan) with UPlanApo (100x) objective (Olympus, Japan) before staining and 2-minute after staining (**Fig. 6A** and **Movie S6**).

### Biofilm assay

The biofilm assay was performed as described previously (46) with some modifications as follows. 0.6 ml or 0.1 ml of liquid culture of the wild type strain or *Δdgc2* strain suspended to an OD730 value of 1.0 was added into 60 mL of BG-11 liquid medium (7.5 mM glucose) in a 100-mL flat-bottom flask or 10 mL of BG-11 in glass plate. Biofilm formation was allowed by incubation at 30°C for 10 days without aeration under low light illumination with a photon flux density of about 5 μmol m^-2^-s^-1^. After incubation, cells attached to the glass bottom and cells remained in the liquid phase, including floc were separately collected. Chlorophyll was extracted from samples incubated in glass plates as follows. For samples remained in the liquid phase, the cell suspension (supernatant) was collected in a 15 mL centrifuge tube by pipetting and centrifugation at 3500 rpm for 5 min at RT to make a pellet. Then, 2 mL of methanol was added to the centrifuge tube (liquid-remained sample) or glass plate (glass-attached sample). The mixture was stirred well and incubated for 5 min at RT. After centrifugation at 10,000 rpm for 10 min at RT, the absorbance of each supernatant at 665 nm was measured to determine the total chlorophyll *a* content (**Fig. 8B, 8C** and **Fig. S4**, 51). The chlorophyll content derived from cells remained in the liquid phase and that from bottom-attached cells were designated as *Cs* and *Cb*, respectively.

### Imaging of cells

Time lapse images for colony patterns on agar plates were taken under the SZX16 microscope (Olympus, Japan) with the SDFPLAPO 0.5XPF (0.5x) objective (Olympus, Japan) using the Moticam Pro282B CCD camera (Motic, China), and Motic Images Advanced 3.2 (Motic, China) was used as the control software (**Figs. 1A, 1D** and **Fig. 7C**). For microscopic observations with higher resolution, we used the IX71 inverted microscope (Olympus, Japan) connected to a cooled CCD camera (Pixis:1024, Princeton Instrument, USA, or Retiga EXi Fast 1394, Qimaging, Canada) and controlled by the SLIDEBOOK 4.2 software (Intelligent Imaging Innovations). For observations shown in **Figs 2 ∼ 4 and 5A, Figs. S2** and **S3, Movies S2 to S5**, the 1.25x objective lens (PlanApo N 1.25x, N.A. 0.04, W.D. 5.1 mm, Olympus, Japan) was used. For observations shown in **Figs. 7D** and **7E, Movie S1**, a 4x objective lens (Uplan FL10×2, N.A. 0.30, W.D.10 mm, Olympus, Japan) was used. The TG-5 digital camera (Olympus, Japan) was used to capture images of cells in glass flasks shown in **Fig. 8A** and cells on plates shown in **Fig. 8D** after biofilm formation. Image J 1.52a (NIH, USA, 52) was used for image processing and analysis. Statistical analysis was performed using the free software R-4.0.3 (64-bit,53). For passing count image shown in **Fig. 5**, the comet-like cluster was tracked manually at the presumed leading position and their positions were recorded. In order to display the trajectory on images, we used a macro in ImageJ, so that the plotted positions from tracking images continuously changed the color from magenta to cyan.

### RNA analysis

For sampling to extract RNA and do RT-qPCR analysis, cells were initially grown in 60 mL of liquid BG-11 media (chloramphenicol and kanamycin were added if necessary) under continuous illumination with white fluorescence lamps of 20 μmol photons m^-2^ s^-1^ for about 7 days, allowing cells to enter the logarithmic growth phase. Cells with a bacterial volume corresponding to the integration of OD_730_ value and the liquid volume (ml) of 5.0 were centrifuged and concentrated to 300 μL. The resulting cell suspension was inoculated onto 120-mL fresh BG-11 medium supplemented with antibiotics and incubated under continuous illumination. Cells at the logarithmic growth phase with OD_730_ of about 1.0 ± 0.4 were collected so that the integration of the OD_730_ value and the liquid volume (ml) would be 3.0. Cells were immediately washed with chilled 50-mL distilled water with each centrifugation at 4°C for 10 min at 3,500 rpm, and suspended in 300 μL of water with four zirconia beads with a diameter of 3 mm (Nikkato, Japan) in 2-ml screw-capped tubes, and stored at -80°C until use. For RNA extraction, the acid hot phenol method (13) was used with some modifications. The stored cells were crushed with the MultiBeads Shocker (Yasui Kikai, Japan) with three cycles of crushing at 2500 rpm for 60 sec followed by 10-sec rest-. For cell lysis, the ISOGEN reagent (Nippon Gene, Japan) was used. RT-qPCR was performed using 1 μg each of total RNA as template for reverse transcription reaction with Super Script III (Thermo Fisher Scientific, USA). 4 μL from the 100-μL cDNA samples were subjected to RT-qPCR with appropriate primer sets shown in **Table S1** and Fast SYBR Green Master Mix (Thermo Fisher Scientific, USA) using the StepOnePlus Real-Time PCR System (Thermo Fisher Scientific, USA). The *ΔΔC*_*T*_ method (54) was used to quantify the expression levels.

### Database search

Sequences for nine putative diguanylate cyclase genes (*LBDG_00200, LBDG_02920, LBDG_10020, LBDG_31460, LBDG_34320, LBDG_34280, LBDG_44250, LBDG_28070* and *LBDG_53130*) were obtained with NCBI or KEGG database. SMART (a Simple Modular Architecture Research Tool, Research Tool (http://smart.embl-heidelberg.de/smart/set_mode.cgi) was mainly used to search for domains. In addition, NCBI’s Conserved Domains (https://www.ncbi.nlm.nih.gov/Structure/cdd/wrpsb.cgi) and EMBL-EBI’s Pfam (https://pfam.xfam.org/) were also employed. We performed a BLASTp search based on Dgc2 in *L. boryana*. While limiting the bacterial phylum, we searched for genes encoding proteins in which CHASE2 and GGDEF domains and the transmembrane region are conserved, and compared the amino acid sequences of nine identified genes and *L. boryana dgc2*. Multiple alignments were generated with SnapGene software (from GSL Biotech; available at snapgene.com) with the Clustal Omega, MAFFT, MUSCLE, or T-coffee tool.

### Resequencing of *L. boryana* E22ml’ genomes and variant calling

Genomic DNA library was constructed using the tagmentation method (Nextera XT, Illumina, San Diego, CA, USA) and was selected with an average insert size of around 400-1000 bp using AMPure XP beads. Sequencing of E22ml’ and E22ml-dg5 were performed with a MiSeq v3 600PE kit (Illumina). Sequencing data were available under accession numbers DRR277765 (E22m1-dg5) and DRR277766(E22m1’), respectively. Breseq (version 0.36; 55) was used to find SNVs and indels and E22m1-dg5 was used to eliminate background mutations in E22m1’. To find large indel and other genomic rearrangement event, SV-Quest v1.0 (https://github.com/kazumaxneo/SV-Quest) was used with default settings.

## Data availability

The strain used in the current study is available at the corresponding author on reasonable request.

## Authors’ contributions

KT and HI conceived and designed the experiments; YF isolated the original E22m1’ mutant; KT, WK, HY, KY, KU and KI performed experiments; KT, YF and HI analyzed the data; KT and HI wrote the manuscript. All authors read and approved the final manuscript.

## Acknowledgements

We thank the members of Iwasaki and Fujita laboratories for their valuable discussion.

## Supplementary information

**Table S1.**
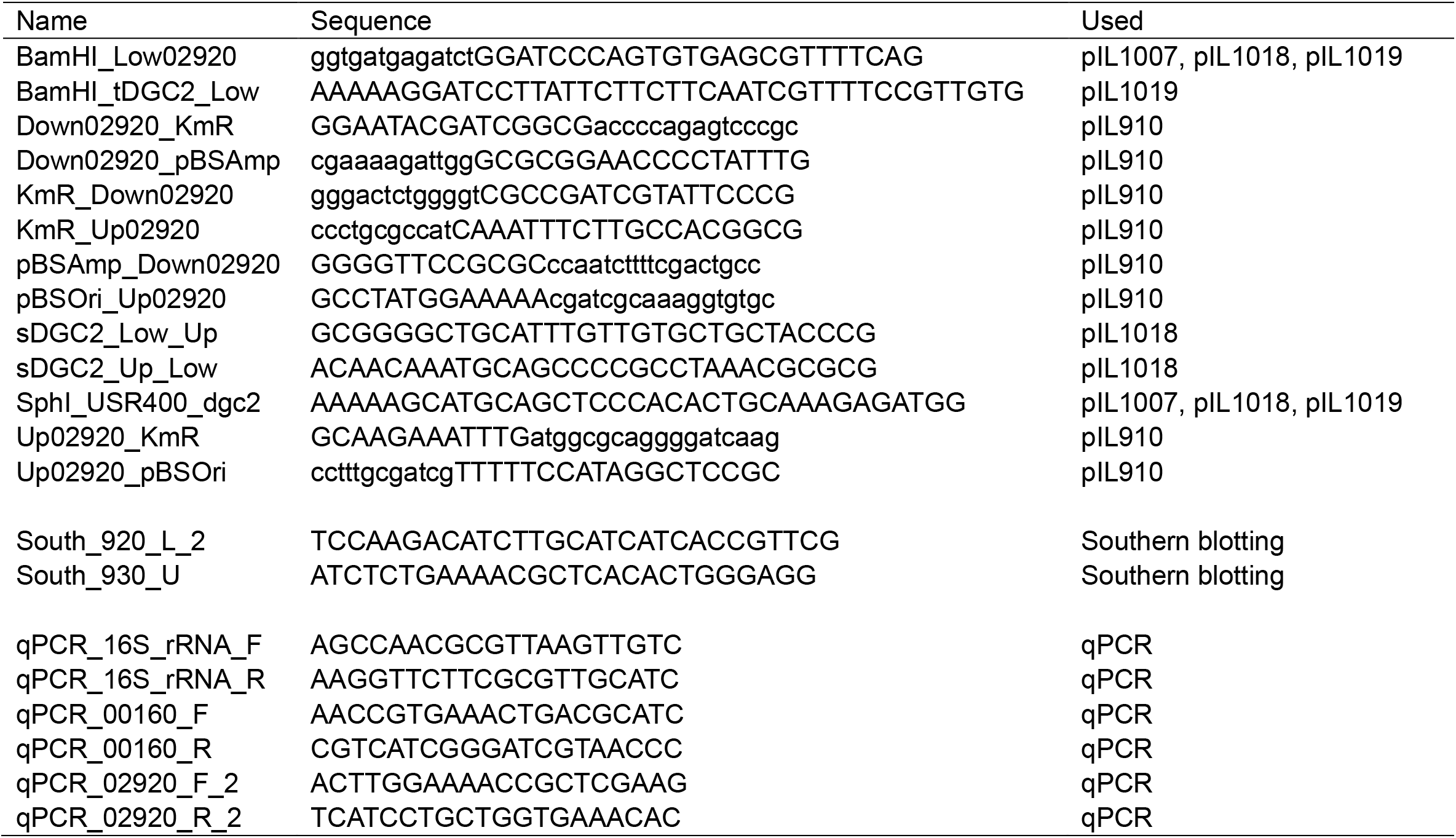
Oligonucleotide primers used in this study.

**Table S2.** Comparison of *dgc2* expression levels.

| comparison | *t*-value | *p*-value |
| --- | --- | --- |
| WT vs $\Delta dgc2$ | 3.3215 | 0.04491 |
| WT vs $\Delta dgc2; dgc2ex$ | -7.0408 | 0.005886 |
| WT vs *dgc2ex* | -1.0479 | 0.3451 |

The expression level of *dgc2* was compared between the wild type and $\Delta dgc2$ , $\Delta dgc2; dgc2ex$ or *dgc2ex* strains using qPCR (*n*=4), respectively. The *t*-values and *p*-values of the Welch's t-test are shown. A significant difference was found between the wild type and the $\Delta dgc2$ or $\Delta dgc2; dgc2ex$ strains (*p*<0.05).

**Table S3.**
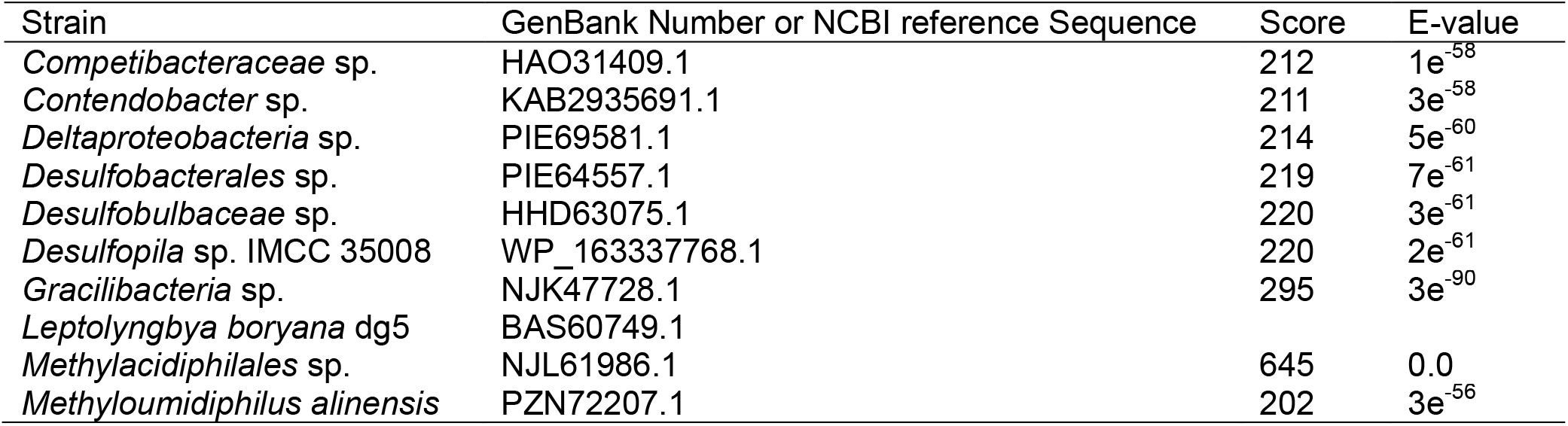
Bacterial strains harboring genes encoding Chase2-containing Dgcs.

**Figure S1. In *L. boryana*, nine genes were found encoding proteins harboring the GGDEF motif**.

We found nine genes encoding proteins harboring the GGDEF motif. We numbered the ORFs in the order in which they were assigned as “LBDG_”, and organized them as Dgc1-Dgc9. Dgc1 has only the REC domain, and Dgc3 had only the REC and EAL domains (gray pentagon) in addition to the GGDEF domain, respectively. EAL domains are also observed in Dgc5 and Dgc7. Dgc5 harbors Transmembrane region, and Dgc7 does two PAS and PAC domains. Dgc4, 6, 8 and 9 are similar. They have REC domain and Trans_reg_C (Transcriptional regulatory protein, C terminal; white rhombus). The Hpt domain is also common and found in three genes other than Dgc6.

**Figure S2. Collective behavior of the original *dgc2***^***−***^ **strain**.

**A**. Time-lapse images of the *dgc2*^*−*^ strain on solid media. Whole images are summarized in **Movie S2** (right). **B**. A kymograph of colonies represented by yellow dashed lines shown in panel **A**. Representation is essentially the same as **Fig. 2**. Magenta arrowhead represents a rotating disk, while cyan and blue arrowheads represent comets.

**Figure S3. Disintegration of rotating disks**.

**A**. Time-lapse images of a disk rotating in the CW direction in the *Δdgc2* strain. Whole images are summarized in **Movie S5B**. Kymograph on the upper part (yellow bar) and the lower part (blue bar) of the disk are shown on the left, as represented in **Fig. 4**. At around 150 min, the edge of the disk began to break up into a comet-like cluster. At around 300 min, lesser amount of cells remained at the center, while its rotation was no longer obvious (see also **Movie S5B**). **B**. Time-course of an exceptional disk cluster rotating in the CCW direction, while the disk disintegrated after collision with some comets. Time-lapse images and three kymographs are shown. Whole time-lapse images are summarized in **Movie S5D**. Color of the three bars on the image at 0 min are correlated with that of kymograph. Red and blue arrowheads represent comets which collided with the disk, while yellow and green arrowheads represent comets which dissociated from the disk. For details, see text.

**Figure S4. Enhanced biofilm formation in the *Δdgc2* strain**.

The amount of chlorophyll extracted from cells attached onto the glass-bottom (biofilm; *Cb*) or cells in liquid phase (*Cs*) in the wild type or *Δdgc2* strain (*n*=20 for both strains). Asterisks indicate significant difference between *Cs* and *Cb* from the wild type strain, and between the *Cb* in the wild type and that in *Δdgc2*strain (Welch’s t-test without assuming equal variances, *p*<0.05).

**Movie S1:** Time-lapse images of colonies in the wild-type (upper left), *dgc2*^*-*^ (upper middle), *Δdgc2* (lower left) and transgenic strains which overexpressing *dgc2* in either the *Δdgc2* (*Δdgc2;dgc2ex*, lower right) strains on solid media for 2 h.

**Movie S2**. Collective behavior of the *Δdgc2* (left) and *dgc2*^*-*^ (right) strains on solid media.

**Movie S3**. Wandering (right) and rotating (right) clusters are shown (extracted from **Movie S2**).

**Movie S4**. Transition of migrating cluster types. **A**. Four comets were unified to a larger one. **B**. Integration of a comet into a disk. **C**. Separation of a comets from a disk.

**Movie S5**. Stable rotation with CCW direction. **A**. A disk continued rotating in CCW direction stably for 300 min. **B**. A CW disk rotating in CW direction emitted comet-like clusters and disintegrated. **C**. This disk had been stable in the CCW direction, but started rotating backwards and disintegrated suddenly. **D**. Rotation of a disk disturbed by collision with comets. The disk collided with other comets at around 185 and 400 minutes, and ones popped out at around 300 and 840 minutes.

**Movie S6**. EPS released from gliding filaments of *Δdgc2*. India ink was dropped on filaments of *Δdgc2* strain. The trajectory of the filament was stained black with the ink.

